# Nuclear-BRITE: an enhanced MS2 system for live-cell imaging of nuclear RNAs

**DOI:** 10.64898/2026.07.07.736975

**Authors:** Adam D. Cawte, Bara Cihlova, Aleksander T. Szczurek, Robert J. Klose, Neil Brockdorff

## Abstract

Analysing the dynamic behaviour of specific RNAs using MS2 stem-loop arrays bound by fluorescently tagged proteins, provides important insights into their biological function. However, high levels of background resulting from the accumulation of unbound stem-loop binding proteins presents a significant limitation to this approach, especially in relation to RNAs that function in the cell nucleus. Here we describe Nuclear-BRITE, an improved MS2-based methodology that significantly reduces nuclear background signal without perturbing the abundance, localisation or function of RNAs of interest. We demonstrate the applicability of this novel approach for fast live-cell single-molecule imaging of endogenously tagged nuclear long non-coding RNAs (Xist, Kcnq1ot1, Malat1), processed mRNAs (Polr2a mRNA), mRNA introns (Hprt and Hspg2), and short functional RNAs (7SK, U1 snRNA, U3 snoRNA, 5S rRNA and scaRNA-8). Accordingly, Nuclear-BRITE provides a valuable new tool to enable live-cell imaging-based analysis of the full range of RNAs that function in the cell nucleus.

## Introduction

RNA molecules perform a wide range of essential functions within the cell nucleus, including the regulation of chromatin and transcriptional output^1–4^, RNA splicing^5–7^, RNA modification^8,9^ and ribosomal biogenesis^10,11^. Messenger RNA (mRNA) is produced by transcription of protein-coding genes^12^. In the nucleus, newly synthesised pre-mRNA undergoes processing, including addition of a 5′ cap, removal of introns by splicing, and addition of a 3′ poly(A) tail. Mature mRNA is then exported to the cytoplasm for translation^13^. Ribosomal RNA (rRNA) is transcribed and processed mainly within the nucleolus, a specialised region of the nucleus. rRNAs associate with imported ribosomal proteins to form the small and large ribosomal subunits, which are subsequently exported to the cytoplasm^10^. Several small nuclear RNAs (snRNAs), including U1, U2, U4, U5 and U6, combine with proteins to form small nuclear ribonucleoproteins (snRNPs). These are major components of the spliceosome, which removes introns from pre-mRNA^7^. Small nucleolar RNAs (snoRNAs) function primarily in the nucleolus. They guide the processing and chemical modification of rRNAs, including methylation and pseudouridylation, helping produce functional ribosomes^14,15^. Small Cajal body-specific RNAs (scaRNAs) are functionally related to snoRNAs, but instead of primarily modifying rRNA, they guide the chemical modification of snRNAs within sub-nuclear compartments known as the Cajal bodies^9^. The nucleus also contains numerous long non-coding RNAs (lncRNAs). These can regulate gene expression through diverse mechanisms, including recruiting proteins to specific genomic regions and influencing chromatin structure^1,3^. A well-known example is Xist, which contributes to X-chromosome inactivation (XCI) in mammals^16,17^.

Defining the properties of RNAs that are localised to the nucleus is central to understanding their function in fundamental biological processes. One approach to address this question is to quantify the dynamic behaviour of RNAs of interest using fluorescent tagging and imaging, for example using the MS2 system^18^. The MS2 system works by incorporating an exogenous array of MS2 bacteriophage RNA stem-loops which are bound by fluorescently labelled MS2 capsid proteins (MCP)^19^. The MS2 system commonly uses a nuclear localisation sequence (NLS) on MCPs, which facilitates the labelling of nascent RNAs produced in the nucleus and additionally enhances imaging conditions in the cytoplasm when visualising mature mRNAs^20,21^. The enrichment and high affinity of fluorescent MCP on MS2 stem-loop arrays enables the detection of single RNA transcripts in the cytosol. However, a major obstacle in visualising nuclear RNAs using this approach is the retention of excess nuclear localised fluorescent protein which hinders the study of nuclear RNAs. Previously described methods aimed at mitigating excess fluorescent capsid protein include the use of orthologous RNA aptamer capsid protein pairings (MS2 and PP7) fused to split fluorescent protein moieties^22^, RNA-stabilised capsid proteins in which excess fluorescent protein is degraded in the absence of RNA^23–25^, and synthetically derived RNA aptamers with high affinity for fluorogenic ligands that circumvent the use of fluorescent proteins^26–34^. However, these approaches have not been successfully applied to endogenously tagged RNAs and instead rely on high transient expression of plasmid encoded RNAs of interest. Endogenous tagging is a critical determinant to ensure RNAs of interest retain physiological expression levels and cellular function.

To overcome the limitations of current methods for imaging nuclear RNAs we have adapted the MS2 system by addition of an ERT2 tag, widely used to regulate availability of DNA interacting enzymes such as Cre recombinase ^35,36^, to ensure excess fluorescent MS2 capsid protein (MCP) preferentially localises to the cytoplasm. A key advantage of basing our methodology around the MS2 system is that it is well established for imaging RNAs at single-molecule resolution and can be used to endogenously tag different RNAs with minimal perturbation of function^37–40^. Additionally, availability of ERT2 tagged proteins is tuneable through addition of Tamoxifen (OHT), providing an option to induce RNA labelling in a controlled manner. We demonstrate that this method, termed Nuclear-background reduced RNA imaging using tamoxifen and ERT2 (Nuc-BRITE) can be used to endogenously tag a broad range of long- and short-RNAs in the cell nucleus with no significant effect on either RNA levels or RNA function. The marked reduction in background levels of fluorescent MCP in the nucleus enables single-molecule resolution and quantitative measurement of the dynamic behaviour of different nuclear RNAs.

## Methods

### Cell Culture

Mouse embryonic stem cells (mESCs) were cultured at 37°C and 5% CO_2_ in ESC media: High Glucose Dulbecco’s Modified Eagle Medium no pyruvate (Life Technologies) supplemented with 10% Foetal Bovine Serum (Sigma), 0.1mM non-essential amino acids, 2mM L-glutamine, 50μM β-mercaptoethanol, 100 U/ml Penicillin/100 μg/ml Streptomycin (Life Technologies) and 1000U/mL Leukemia Inhibitory Factor (LIF) conditioned medium (made in-house). mESCs were grown on gelatinised plates, on mitotically arrested mouse SNLP feeder cells (STO mouse fibroblasts expressing Neomycin, Puromycin resistance and LIF genes). Stocks of feeders were frozen down from mitotically arrested SNLPs cultured in mitomycin-C (Sigma Aldrich) at 1µg/ml for 2h.

### Generation of Nuclear-BRITE cell lines

To initially test the enhancement and behaviour of different Nuc-BRITE imaging constructs, we used a previously developed cell line that had an MS2v7 x24 array inserted into the doxycycline inducible Xist gene (129 allele) in a female Castaneus/129 interspecific mESC cell line (iXist-ChrX^129^–^MS2v7^)^41^. The different pCAG driven imaging constructs (Nuc-BRITE-tdStayGold, Nuc-BRITE-Halo and NLS-tdMCP-tdStayGold) were stably integrated into the iXist-ChrX^129^–^MS2v7^ cell line by transfecting 1 µg of plasmid DNA with lipofectamine 3000 (Thermo Fisher) following the manufacturers protocol into cells seeded on a 6-well plate. Positively transfected cells were expanded up to 10 cm dishes and selected for using Neomycin at 400 µg/ml for ∼14 days. Individual mESC clones were isolated via serial dilution and seeding on 96-well plates which were later assessed for efficient transgene expression via imaging.

For later experiments, we opted for stable Rosa26 expression to minimise variation in Nuc-BRITE expression levels. For this we used the female Castaneus/129 interspecific iXist-ChrX^129^ mESC cell line^42^ as the parental cell line for Rosa26 integration. CRISPR/Cas9 homology directed repair (HDR) was used to endogenously modify the iXist-ChrX^129^ mESCs to generate various Nuc-BRITE imaging cell lines. First, the Rosa26 locus on Chr 6 was targeted with a Rosa26 specific sgRNA and Rosa26-AdSA-tdMCP-tdStayGold-ERT2 (Nuc-BRITE) homology plasmid. Cells were transfected in 6-well plates using lipofectamine 3000 (Thermo Fisher) with 1 µg of sgRNA plasmids (based around pX459-v2.0 puro, addgene plasmid #62988) and a 6:1 molar ratio of homology plasmid. Following 24h of plasmid expression, cells were split and seeded into three densities, 1:2, 1:3 and 1:6 on 10cm dishes. After a further 24h incubation, puromycin was used for selection at a concentration of 3.5 µg/ml for a further 48h. Following puromycin selection, cells were recovered for ∼ 10 – 14 days at which time single colonies were picked and seeded into 96-well plates. Positive Rosa26-Nuc-BRITE cell lines were screened by live-cell imaging and PCR (PhantaFlash – Vazyme) to confirm an even fluorescent signal and Rosa26 integration. The selected clones were additionally tested for puromycin sensitivity to enable further rounds of CRISPR/Cas9 engineering.

### Generating MS2v7 x24 tagged cell lines

An MS2v7 x24 array (addgene plasmid # 140705) was cloned into plasmids with homology for Polr2a mRNA, Hprt intron 1, Hspg2 intron 33, Malat-1, Xist and Kcnq1ot1. For Polr2a mRNA, Hprt intron 1, Malat-1 and Kcnq1ot1, the homology regions (HR) and MS2v7 x24 arrays were cloned into a pBlueScript II backbone. PCR of HRs (∼500bp – 1kb) and subsequent restriction enzyme digest (NEB) of both HRs and MS2v7 x24 from pUC-MS2v7 x24 was used to generate the inserts for combined ligation (T4 DNA ligase, ThermoFisher) into a calf intestinal phosphatase treated (NEB) and linearised pBlueScript II backbone. For Xist and Hspg2 intron 33, Gibson assembly (ClonExpress II, Vazyme) was used to combine PCR products for each HRs and MS2v7 x24 array into a minimal plasmid backbone (pHR). MS2v7 x24 tagged cell lines were generated as previously described above for Rosa26-Nuc-BRITE, using site specific CRISPR/Cas9 HDR and puromycin selection. Positive clones were selected based on PCR screening for correct MS2v7 x24 insertion and validated via live-cell imaging. A list of CRISPR sgRNAs, cloning and screening primers for MS2v7 x24 constructs can be found in Supplementary Table 1.

### Generating internally tagged short ncRNA cell lines

To incorporate MS2 into 7SK, U1a1 snRNA and U3a snoRNA, PCR products of HRs incorporating single MS2 stem-loops were amplified from genomic DNA and cloned together using Gibson assembly into pHR. MS2 stem-loop insertion was chosen based on internal stem-loops (SL) within each RNA which are predicted, avoiding known essential elements, for example those bound by key RNA binding proteins. For 7SK, U1a1 snRNA, U3a snoRNA and murine scaRNA-8 we designed constructs to tag SL1, SL4, SL2 and SL2 in each RNA respectively. For 5S rRNA, an MS2 x3 array was incorporated at the 3’ end of the RNA similar to that published previously^26,27^. The MS2 x3 array was amplified from the MS2v7 x24 array, incorporated into pSLQ-5S-F30-control, addgene plasmid #127582, by replacing the F30 scaffold. Following this, the mU6-5S-MS2 x3 sequence was amplified by PCR along with HRs of a unique site adjacent to the 5S rRNA cluster on Chr 8 for Gibson assembly into pHR. For scaRNA-8, the RNA sequence was amplified from genomic DNA to incorporate an internal MS2 SL and this sequence was incorporated into pSLQ-mU6-Puro plasmid (adapted from addgene plasmid #127582) using Gibson assembly. This plasmid was used to generate randomly integrated mESC clones following puromycin selection. Short ncRNAs that were endogenously targeted included 7SK, U1a1 snRNA, U3a snoRNA and 5S-MS2 x3 rRNA. Site specific CRISPR/Cas9 HDR in mESCs, as previously described, was used to introduce MS2 tagged RNAs. All clones except U1a1 snRNA generated endogenously tagged ncRNAs that were efficiently expressed. The U1a1 snRNA clone was initially selected for its stable expression of U1a1-MS2 as visualised via live-cell imaging, however, further analysis showed it was stably integrated in the genome away from the endogenous U1a1 site on Chr11. A list of CRISPR sgRNAs, cloning and screening primers for short ncRNA constructs can be found in Supplementary Table 2.

### qPCR

Quantitative PCR was carried out on wild-type, Nuc-BRITE, Polr2a-MS2v7×24, and Malat-1-MS2v7×24 cell lines to determine the effect Nuc-BRITE and MS2 incorporation had on RNA abundance. RNA was isolated using Trizol reagent following the manufacturers protocol (ThermoFisher). Amplification of cDNA used SuperScript III reverse transcriptase (ThermoFisher), random primers (100 ng total) and 1 µg of isolated RNA in a total reaction volume of 20 µl following the manufacturers protocol. The Polr2a forward primer spanned the junction between exon 3 and 4 and was used in conjunction with a reverse primer further downstream in exon 4 to amplify mature RNA products. The 5’ end of Malat-1 (170 bp) was amplified to assess its levels in the different lines. Gapdh expression was used to normalise the Ct values for each sample in comparison to wild-type cells and the variation in each sample was taken from 4 technical replicates. A Rotogene qPCR machine and Taq Pro SYBR qPCR master mix was used for amplification (Vazyme). A threshold of 0.01 was used to calculate Ct values at the start of the log phase of PCR amplification. Data was plotted in Igor Pro 9 (Wavemetrics). All qPCR oligos can be found in Supplementary Table 3

### RNA-FISH and immunofluorescence

mESCs were grown on gelatin-coated No. 1.5H precision coverslips (170 ± 5 µm; Marienfeld Superior) in 6-well plates. Coverslips were washed with PBS and fixed with 3.7% formaldehyde in PBS for 10 min at room temperature. Cells were rinsed three times with PBST.5 (PBS + Tween20 0.05%), permeabilized with 0.2% Triton X-100 in PBS for 10 min, washed again three times with PBST.5, followed by a 30 min incubation in 70% EtOH at room temperature. Coverslips were then dehydrated with subsequent washes with 80%, 90% and 100% EtOH for 5 min each and briefly dried.

FISH probes were generated from either a 1.5kb PCR product of the Polr2a exon 29 / 3’UTR sequence or a 6kb PCR product of Malat-1. Nick translation (Enzo) of the PCR products was used to incorporate Cy5-dUTP (Enzo) following the manufacturers protocol with incubation for 2 h at 15°C. Cy5 labelled probes, 5 µl per hybridization, were ethanol precipitated with 1/3 volume of 10 mg/ml salmon sperm DNA, 1/10 volume of 3 M NaOAc and 3 volumes of 100% EtOH. After spinning and washing with 70% EtOH, the pellet was air dried and resuspended in deionized formamide (Sigma). Probes resuspended in formamide were added to the hybridisation mixture to a final concentration of 2xSSC buffer, 50% Formamide (containing probes), 5% Dextran sulphate, 1 mg/ml BSA, 2 mM Vanadyl Ribonucleoside Complex.

Coverslips were hybridized with 30 µl probe/hybridization buffer mix that was denatured at 75°C for 5 min before briefly being chilled on ice. Hybridization was carried out in a humid chamber overnight at 37°C. The next day, coverslips were washed three times with prewarmed 50% formamide/2x SSC at 42°C for 5 min each, and subsequently three times with 2x SSC at 42°C for 5 min each. Coverslips were rinsed with water to remove excess salts, stained with DAPI (1 µg/ml -Sigma) and mounted with Everbrite (biotium) mounting medium to improve Cy5 fluorescence stability, on Superfrost Plus microscopy slides (VWR). Slides were dried, sealed using clear nail polish, and cleaned for imaging.

For immunofluorescence, cells were cultured on gelatin-coated No. 1.5H precision coverslips (170 ± 5 µm; Marienfeld Superior). Coverslips were washed once with PBS prior to fixation in 3.7% formaldehyde for 10 mins. Coverslips were washed 3x with PBST.5, permeabilized with 0.2% Triton X-100 in PBS for 10 min and washed again 3x with PBST.5. Coverslips were blocked in 4% BSA diluted in PBST.5 for 30 mins. A primary antibody against Fibrillarin (Abcam ab5821) was used at a 1:200 dilution in 4% BSA and incubated overnight at 4°C. Coverslips were washed 3x in PBST.5 and incubated with an AlexaFluor647 conjugated goat anti-rabbit IgG secondary antibody (Invitrogen −1:2000) for 1h in 4% BSA. Coverslips were washed 3x in PBST.5 followed by washes in PBS and stained with DAPI (1µg/ml -Sigma) prior to mounting in Everbrite (biotium).

### Live-cell imaging

For imaging experiments, mESCs were grown in gelatinised ibidi 8-well chamber slides (ibidi-#80806) in the absence of feeders and imaged in live-cell imaging media: FluoroBrite DMEM supplemented with 20 mM HEPES, 10% Foetal Bovine Serum, 0.1mM non-essential amino acids, 2mM L-glutamine, 50μM β-mercaptoethanol, 100 U/ml Penicillin/100 μg/ml Streptomycin (Life Technologies) and 1000U/mL Leukemia Inhibitory Factor (LIF) conditioned medium. To induce Xist expression in iXist-ChrX^129^–^MS2v7^ mESCs, doxycycline (Sigma) was used at 1 µg/ml for 24h. For Nuc-BRITE-tdStayGold cells, in standard imaging conditions, no tamoxifen was added for RNA labelling. For comparison of Xist RNA in Nuc-BRITE-tdStayGold and Nuc-BRITE-Halo cells, 100 nM tamoxifen (OHT) was added to the media prior to imaging to compare induction of RNA labelling. For 7SK cells treated with NVP-2 (Tocris Bioscience), 500 nM NVP-2 was used for a 30 min treatment prior to imaging.

### Microscopy

For fast frame rate live-cell super-resolution imaging, Nuc-BRITE samples were imaged using a Visitech iSIM super resolution unit built around an Olympus IX83 microscope. The iSIM has 4 diode lasers 405, 488, 561 and 640 nm (10 mW each) that run through the iSIM super-resolution confocal array scanner with images collected on a dual camera set-up housing two Hamamatsu Orca Quest cameras. Fluorescence was filtered using specific pairs of filters for each pair of channels: 488/561, 488/640, 405/640. Fast super-resolution dual-colour imaging used a 1.5 N.A. 100X PlanApo objective, 63 ms exposure and 80% laser power for 200 frames. The dual-camera set-up was aligned across all 4 channels using X, Y and Z tilt adjustments and a TetraSpek bead slide (Thermo Fisher) to compensate for chromatic aberrations.

Long-term live-cell imaging, single time point maximum projections and fixed RNA FISH or IF samples were imaged using an Olympus IX83 widefield microscope using CooLED illumination at 365, 490, 580 and 635 nm through a 1.35 N.A. 60X PlanApo objective, filtered using fixed filter sets (DAPI, GFP, TRITC and Cy5) and images were collected on a Prime 95B sCMOS camera (Photometrics). Single time point images for live-and fixed-cell were acquired as Z-stacks using 10 ms (DAPI) or 200 ms (GFP/TRITC/Cy5) exposure time and 5% (DAPI) and 30% (GFP/TRITC/Cy5) illumination. Long-term imaging of Hspg2 intron splicing acquired 15 µm Z-stacks over ∼1 hour (1 min frame rate) or 8 hours (4 min frame rate) using 70 ms exposure 350 nm per Z-slice and 8% 490 nm (GFP) excitation.

Sing;e particle tracking (SPT) of 7SK-MS2 was imaged using an Oxford Nanoimager (ONI) set up with a 488 nm laser and TIRF objective set at an angle of 45° in HiLO illumination. To image single particles of 7SK-MS2, cells were first imaged with 20% laser for ∼ 250 frames to define the boundary of the nucleus and nuclear specific signal. Following this, cells were bleached using 60% laser power over ∼ 250 frames until single particles could clearly be observed. After this, the laser power was reduced to 20% and an SPT analysis window was used from frame 1000 – 5000. Cells were imaged using 15 ms exposure.

### Image analysis

Analysis of Xist and Polr2a foci intensities or dynamics was undertaken using ImageJ, the Trackmate plugin^43^ and custom MATLAB scripts. In short, cells were cropped, batch processed using the Trackmate batcher plugin which exported trajectory and foci information which was subsequently imported into MATLAB for quantification of foci intensities and diffusive behaviours. More specifically, to quantify intensities and trajectories of Polr2a molecules, cropped images were batch processed with the following Trackmate settings to detect single particles over time and reduce false positives from neighbouring or out-of-focus foci: Laplacian of Gaussian detector, Foci diameter – 300 nm, Quality – 5, Contrast – 0.02, LAP tracker – Frame linking 500 nm, Gap closing 1000 nm, Number of spots in track – 5.

For Xist molecules the Trackmate settings were: Laplacian of Gaussian detector, Foci diameter – 200 nm, Quality – 20, Contrast – 0.02, LAP tracker – Frame linking 200 nm, Gap closing 200 nm, Number of spots in track – 5. Only spots that were allocated to trajectories were analysed and .csv files were imported into MATLAB for quantification and foci intensities were plotted as histograms in Igor Pro 9 (Wavemetrics).

For quantification of diffusive behaviour, images were batched processed as described above and apparent diffusion coefficients were calculated using the msdanalyser suite of processes in MATLAB^44^. Briefly, simple .xml files were exported and subsequently imported as concatenated cell arrays into MATLAB. Cell arrays were then processed individually in msdanalyser on a cell-by-cell basis. MSD values were calculated for each trajectory and apparent diffusion coefficients (D*) were calculated using equation 1. and plotted as log_10_D* values in Igor Pro 9.

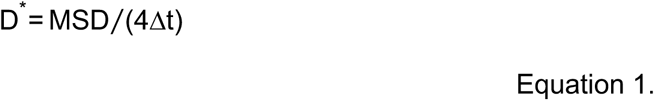

For SPT analysis of Nuc-BRITE, 7SK and 7SK + NVP-2 cells, individual nuclei from each condition were cropped and batch processed using TrackMate and analysed using Laplacian of Gaussian detector, Foci diameter – 300 nm, Quality – 50, NND tracker 600 nm and Number of spots in track – 4. Analysis of trajectories assessed each trajectories apparent diffusion coefficient as described above. Apparent diffusion coefficients were similarly plotted as log_10_D* histograms using Igor Pro 9.

All MATLAB scripts can be found at the following github page: https://github.com/adamcawte/NucBRITE_MATLAB_Code.git

## Results

### Application of Nuclear-BRITE for enhanced imaging of Xist lncRNA

To address the limitations of the MS2 system for endogenous tagging and imaging of RNA in the nucleus, we investigated including an ERT2 tag fused to a tandem dimer MS2 capsid protein (tdMCP) and either a tandem dimer StayGold (tdStayGold) fluorescent protein^45^ or Halo-Tag protein to generate two different Nuc-BRITE constructs. The ERT2 tag is commonly used to retain DNA interacting enzymes in the cytosol. Addition of the estrogen antagonist Tamoxifen (OHT) allows for tuneable nuclear localisation in a concentration dependent manner^35,36^. To test the two Nuc-BRITE constructs we first analysed the long non-coding RNA Xist, making use of an XX mouse embryonic stem cell (mESC) line^42^ into which an MS2v7 x24 array^46^ had been inserted into a doxycycline inducible Xist gene (iXist-ChrX^129^–^MS2v7^)^41^ (Fig. 1a). We then stably integrated constructs encoding the Nuc-BRITE fusion proteins (Fig. 1b and c). Initial experiments demonstrated that Nuc-BRITE-tdStayGold confers enhanced low background imaging of Xist RNA in a OHT independent manner whereas the Halo-Tag fusion protein labels Xist only in the presence of OHT (Fig. 1b and c). The basis for this difference is not known but may reflect different affinities of the two fusion proteins for binding HSP90 which retain ERT2 fusion proteins in the cytosol. Regardless, both systems provide markedly improved labelling of Xist RNA in the cell nucleus. A comparison with conventional MS2 imaging of Xist RNA using a tdMCP-tdStayGold fusion protein with a nuclear localisation sequence (NLS) shows that Nuc-BRITE enhances the ability to observe nuclear Xist foci (Fig. 1d and e). In single-molecule tracking analysis Nuc-BRITE increases signal-to-noise ratios of detected nuclear foci as well as the number and duration of tracks that can be observed per cell (Fig. 1f – h). An extended analysis of Nuc-BRITE tagged Xist RNA, including the demonstration that gene silencing in XCI is unperturbed, is provided elsewhere^41^.

**Fig. 1.**
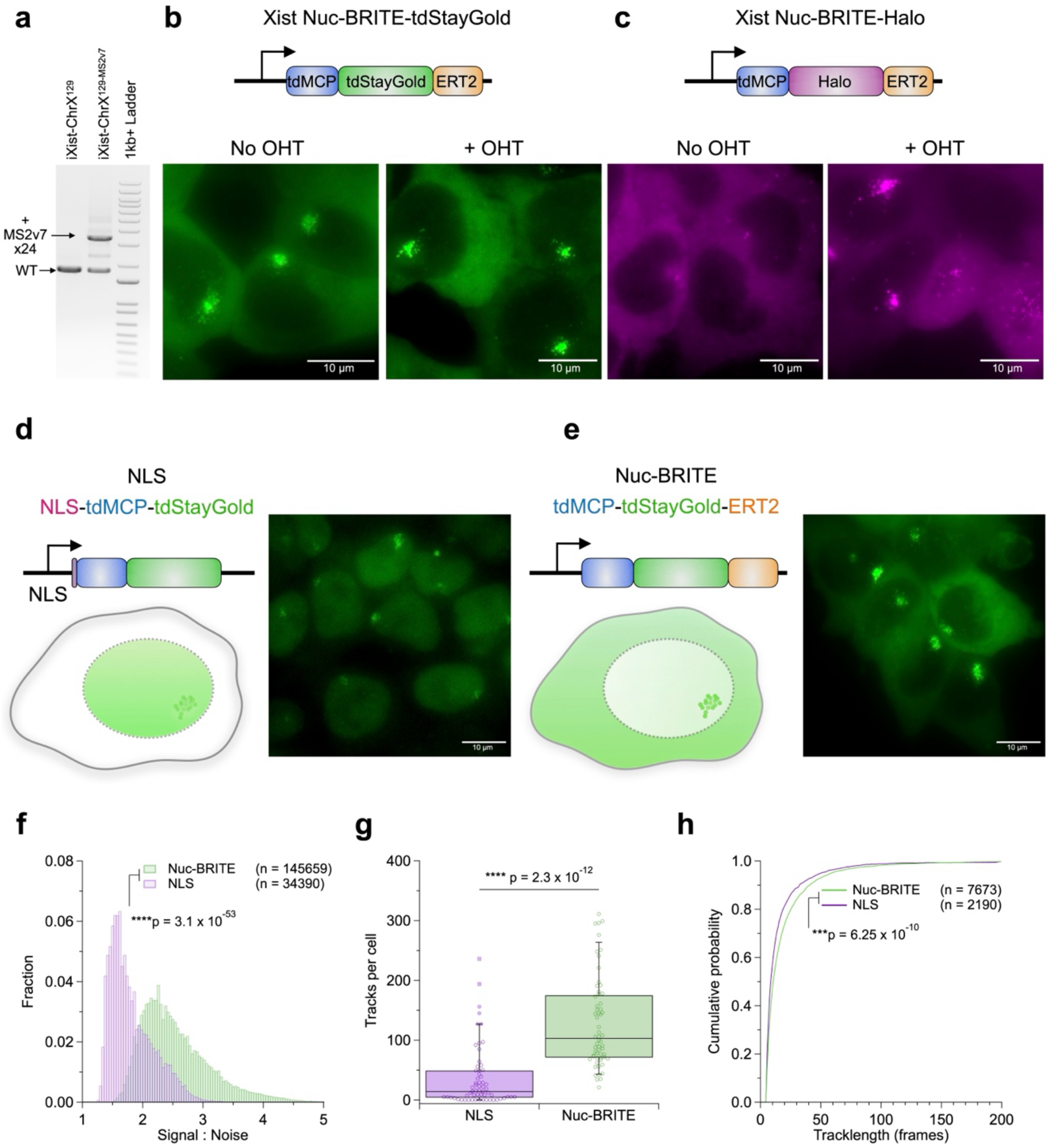
Enhanced imaging of Xist lncRNA with Nuclear-BRITE. a) PCR amplification of genomic sequences across MS2v7 x24 insertion site (Exon 7) in Xist for iXist-ChrX^129^ wildtype and iXist-ChrX^129^–^MS2v7^ cells. b) Schematic illustrating the Nuc-BRITE-tdStayGold construct used to image Xist-MS2v7 (tdMCP – blue, tdStayGold – green, ERT2 – orange) and representative images of dox induced (24h) Xist-Nuc-BRITE-tdStayGold cells in the presence or absence of Tamoxifen (OHT). c) Schematic illustrating the Nuc-BRITE-Halo construct used to image Xist-MS2v7 (tdMCP – blue, Halo – purple, ERT2 – orange) and representative images of dox induced (24h) Xist-Nuc-BRITE-Halo cells stained with Halo-JF549 (100 nM) in the presence or absence of Tamoxifen (OHT). Schematics and representative images of NLS (d) and Nuc-BRITE (e) imaging of Xist in iXist-ChrX^129^–^MS2v7^ cells after 24h dox induction. f) Histograms of signal-to-noise ratios for detected foci in NLS and Nuc-BRITE cells. Number of foci quantified shown as n. g) Box plot of the number of tracks per cell for NLS and Nuc-BRITE cell lines. h) Cumulative probability of track length for NLS and Nuc-BRITE cell lines, number of trajectories analysed shown as n. All data for (f – h) taken from 61 and 60 cells for NLS and Nuc-BRITE respectively. All statistical analysis used single tailed t-tests between distribution of single-molecule intensities or trajectories.

### Nuclear-BRITE is a versatile tool to study a broad range of nuclear RNAs

Encouraged by the results obtained imaging Xist lncRNA, we aimed to investigate if Nuc-BRITE is equally applicable to other endogenously tagged nuclear RNAs. Due to its constitutive fluorescence and ease of use, we went on to use Nuc-BRITE with tdStayGold to analyse different nuclear RNAs of interest. For these experiments we stably integrated Nuc-BRITE into the Rosa26 locus^47^ to ensure even levels of fluorescent protein expression across the different mESC clones. In the absence of MS2 RNA, the majority of Nuc-BRITE signal is cytoplasmic with a small proportion in the nucleoplasm (Supplementary Video. 1). We next tested the ability of Nuc-BRITE to image coding and non-coding RNAs (ncRNAs) at single-molecule resolution in living cells. We endogenously tagged Polr2a mRNA, Hprt intron 1, and the lncRNAs Malat-1, Xist and Kcnq1ot1 with the MS2v7 x24 array^46^ (Fig. 2a and b). Tagging occurred on both alleles in all cases except for Xist and Kcnq1ot1 in which the tag was integrated into an expressed allele (Supplementary Fig. S1 a and b). For all RNAs imaged using Nuc-BRITE, the observed localisation pattern, dynamic behaviour and apparent abundance were in line with expectations (Fig. 1b). Additionally, in cases such as nascent transcript imaging, we were able to clearly observe splicing of the MS2 labelled intron (Hprt intron 1 -Supplementary Video. 1).

**Fig. 2.**
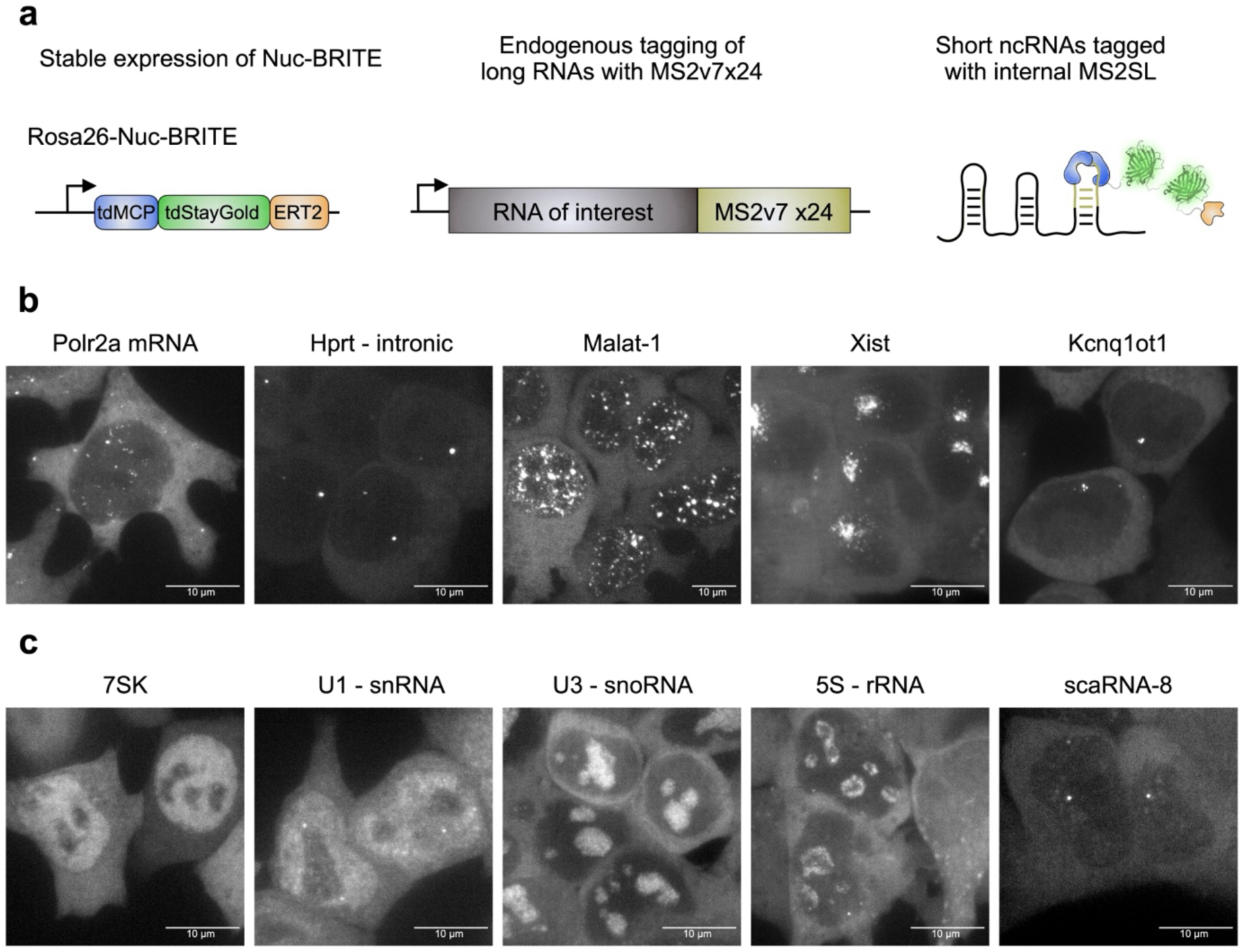
Endogenous tagging of various nuclear RNAs using Nuclear-BRITE. a) Schematics of Nuclear-BRITE fluorescent capsid construct (left), and endogenous RNA tagging with either an MS2v7 x24 array for long RNAs (centre) or an internal MS2 stem-loop for short RNAs (right). B) Maximum intensity projections for long RNAs tagged with Nuc-BRITE – Polr2a mRNA, Hprt intron 1, Malat-1 lncRNA, Xist lncRNA and Kcnq1ot1 lncRNA. C) Maximum intensity projections for short RNAs tagged with Nuc-BRITE – 7SK, U1a1 snRNA, U3a snoRNA, 5S rRNA and murine scaRNA-8.

To assess whether Nuc-BRITE had an impact on RNA stability, we quantified the levels of Polr2a mRNA and Malat-1 in wild-type (WT), Nuc-BRITE alone and MS2v7 x24 tagged cell lines and observed no significant changes in RNA expression levels between tagged and WT cells (Supplementary Fig. S1c). Furthermore, imaging RNA FISH probes against Polr2a mRNA and Malat-1 demonstrated colocalization with Nuc-BRITE signals, demonstrating the specificity of Nuc-BRITE and the retention of localisation patterns seen in unmodified cells (Supplementary Fig. S1d). Comparison of these cell lines highlights how Nuc-BRITE efficiently and specifically labels diverse nuclear RNAs with a wide range of expression levels, over ∼ 3 orders of magnitude between Kcnq1ot1 (∼ 5 molecules per cell) and Malat-1 (∼ 5,000 molecules per cell).

In further experiments we used Nuc-BRITE to tag small nuclear RNAs with a single MS2 aptamer inserted into structural stem-loops of the RNAs^27,48^ (Fig. 2a and c). We endogenously incorporated MS2 tagged 7SK, U1 snRNA, U3 snoRNA, 5S rRNA and murine scaRNA-8 (Supplementary Fig. S2a). As most short ncRNAs have multiple isoforms, we applied different strategies of MS2 incorporation either at a single endogenous locus (7SK -*Rn7sk* and U3 snoRNA -*Rnu3a*), a locus adjacent to a ncRNA cluster (5S rRNA), or stable expression of a randomly integrated ncRNA construct (U1 snRNA and scaRNA-8). Of note, the unique locus expressing 7SK RNA was homozygously tagged as shown by a shift of ∼ 20 bp for full-length 7SK (Supplementary Fig. S2b). All endogenously expressed ncRNAs were observed to have the expected RNA localization pattern, expression level and behaviour in live cells, including the nucleolar import of 5S rRNA during ribosomal biogenesis (Supplementary Video. 2). Additionally, each short ncRNA localised to the correct nuclear body, as confirmed with immunofluorescence against Fibrillarin which associates with both nucleoli and Cajal bodies (Supplementary Fig. S2c). Importantly, labelling of highly abundant ncRNAs such as 7SK, U1 snRNA and U3 snoRNA was evident despite the presumed limited nuclear pool of Nuc-BRITE fusion protein. Similarly, Nuc-BRITE levels in the nucleus remained low with minimally expressed ncRNAs such as scaRNA-8. This shows Nuc-BRITE efficiently labels short ncRNAs at varying levels of abundance, whilst retaining improved nuclear background levels.

### Functional validation of Nuclear-BRITE tagged RNAs

To test whether Nuc-BRITE tagged RNAs retain functions and properties defined in prior studies, we investigated homozygously tagged Polr2a mRNA and 7SK RNAs following puromycin or CDK9 inhibitor (NVP-2) treatment respectively^49,50^. Nuc-BRITE Polr2a mRNAs were observed to form clusters of ∼ 2 – 3 molecules in the cytoplasm. These accumulations have been shown to be associated with active translation and are thus sensitive to puromycin^49^. Qualitative analysis indicated that these larger cytoplasmic foci are reduced in ∼ 2 minutes after addition of puromycin (Fig. 3a and Supplementary Video. 3). In line with this observation, quantitative analysis demonstrated a marked decrease in mean foci intensities over this time course (Fig. 3b). Additionally, quantifying Polr2a mRNA foci in the absence or presence of puromycin shows that the Polr2a mRNA foci detected have reduced intensity and increased mobility in the presence of puromycin (Fig. 3c and d, Supplementary Video. 3).

**Fig. 3.**
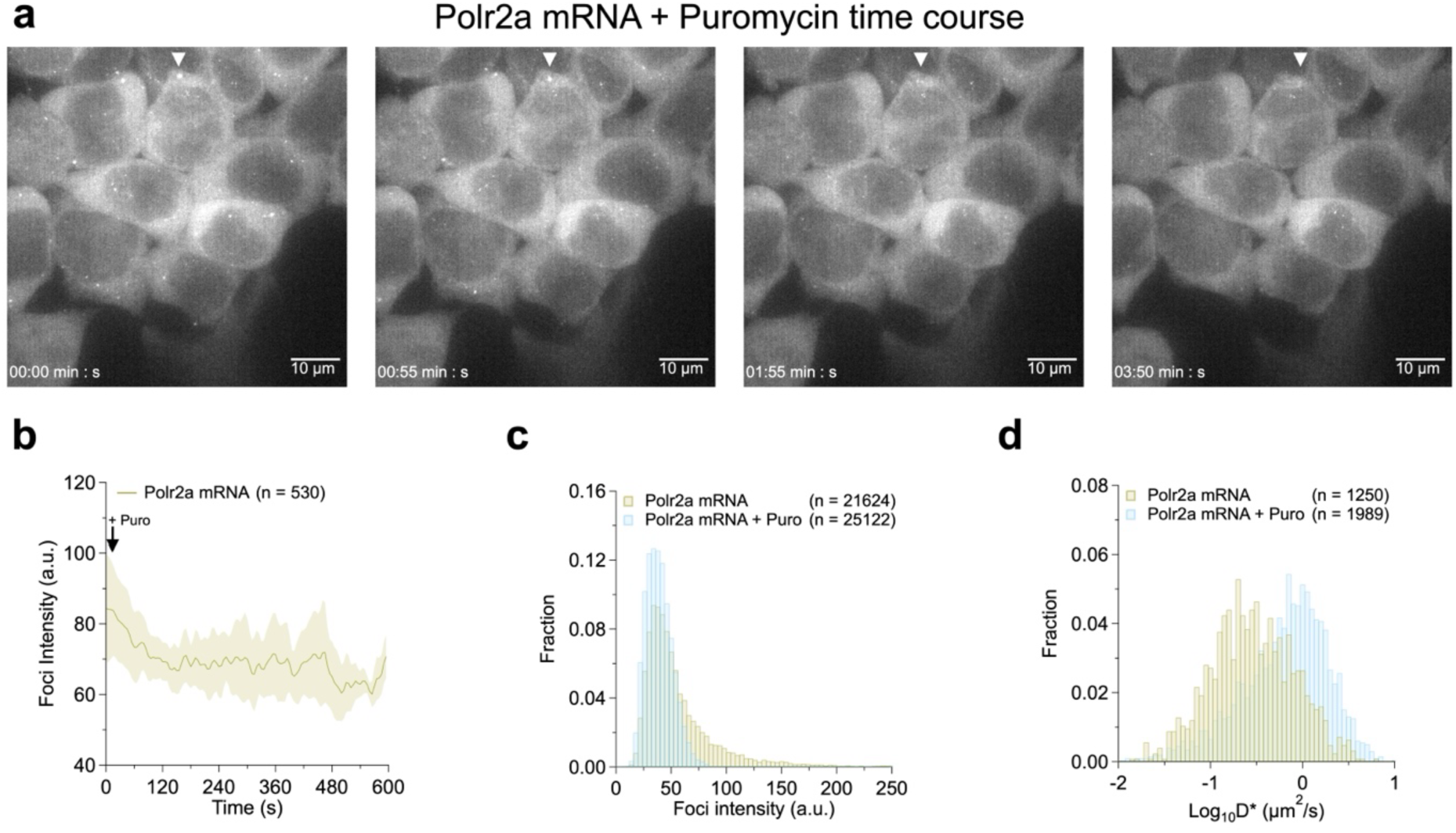
Nuclear-BRITE tagged Polr2a mRNAs are sensitive to puromycin. a) Maximum intensity projection of Polr2a mRNA foci following puromycin addition. Arrow indicates a multimeric Polr2a mRNA focus dissociating over time. b) Mean intensity profile over time for Polr2a mRNA foci following puromycin addition (arrow indicates puromycin addition). Mean shown as solid line and standard deviation depicted as shaded area around the mean. n represents number of single particle trajectories analysed from a total of 22 cells. c) Foci intensity histograms and d) Log_10_D* histograms for Polr2a mRNA foci in the presence (blue) and absence (yellow) of puromycin (2 µM for 1 h). n represents number of foci or trajectories analysed from a total of 39 and 48 cells from Polr2a mRNA and Polr2a mRNA + puro respectively.

A prior study used single particle tracking (SPT) of 7SK associated proteins to demonstrate dissociation of the P-TEFb:7SK snRNP complex and accumulation of various 7SK:hnRNP complexes following CDK9 inhibition with NVP-2^50^. As Nuc-BRITE utilizes tdStayGold, which is as bright and stable as organic Halo dyes^45^, we hypothesised that single molecules of 7SK RNA could be visualised under appropriate imaging conditions for SPT^51^, enabling direct measurement of the dynamic behaviour of 7SK RNA. Thus, we imaged Nuc-BRITE cells lacking MS2 tagged RNA as well as 7SK-MS2 cells in the presence or absence of NVP-2 (Supplementary Video. 4). Nuclear signal for Nuc-BRITE alone and 7SK-MS2 was observed as expected under Highly-inclined Laminated Optical sheet (HiLO) illumination (Fig. 4a and b). After initial imaging to define the nuclear boundary, the samples were subjected to bleaching until an appropriate density of single molecules was observed (Fig. 4c and Supplementary Video. 4). A greater number of trajectories were observed in 7SK-MS2 cells compared to the Nuc-BRITE control sample under the same imaging conditions (Fig. 4d and e). Additionally, the diffusive behaviour of 7SK-MS2 trajectories, as quantified as apparent diffusion coefficients (D*), was distinct from trajectories quantified from Nuc-BRITE cells and accurately ranged from ∼ 0.01 – 10 µm^2^/s, a similar range to that observed for 7SK associated proteins^50,52^ (Fig. 4f). Accordingly, we infer that the smaller less diffusive fraction (∼0.01 – 0.1 µm^2^/s) is linked to 7SK RNP’s engagement with chromatin and subsequent release of active components within the P-TEF-b:7SK RNP complex. The larger more diffusive population (∼ 0.5 – 10 µm^2^/s) on the other hand, likely represents the freely diffusive P-TEF-b:7SK RNP complex. Short incubations with NVP-2 (30 mins) led to a reduction in the slower moving fraction with D* values of ∼ 0.01 – 0.1 µm^2^/s and a modest increase in diffusive molecules with D* values of ∼ 3 −4 µm^2^/s. These data are in line with previous studies which concluded that CDK9 inhibition with NVP-2 reduces 7SK RNP chromatin interactions, triggers P-TEF-b dissociation and enables hnRNP association with the 7SK RNP, in turn leading to modest increases in diffusive behavior^50^.

**Fig. 4.**
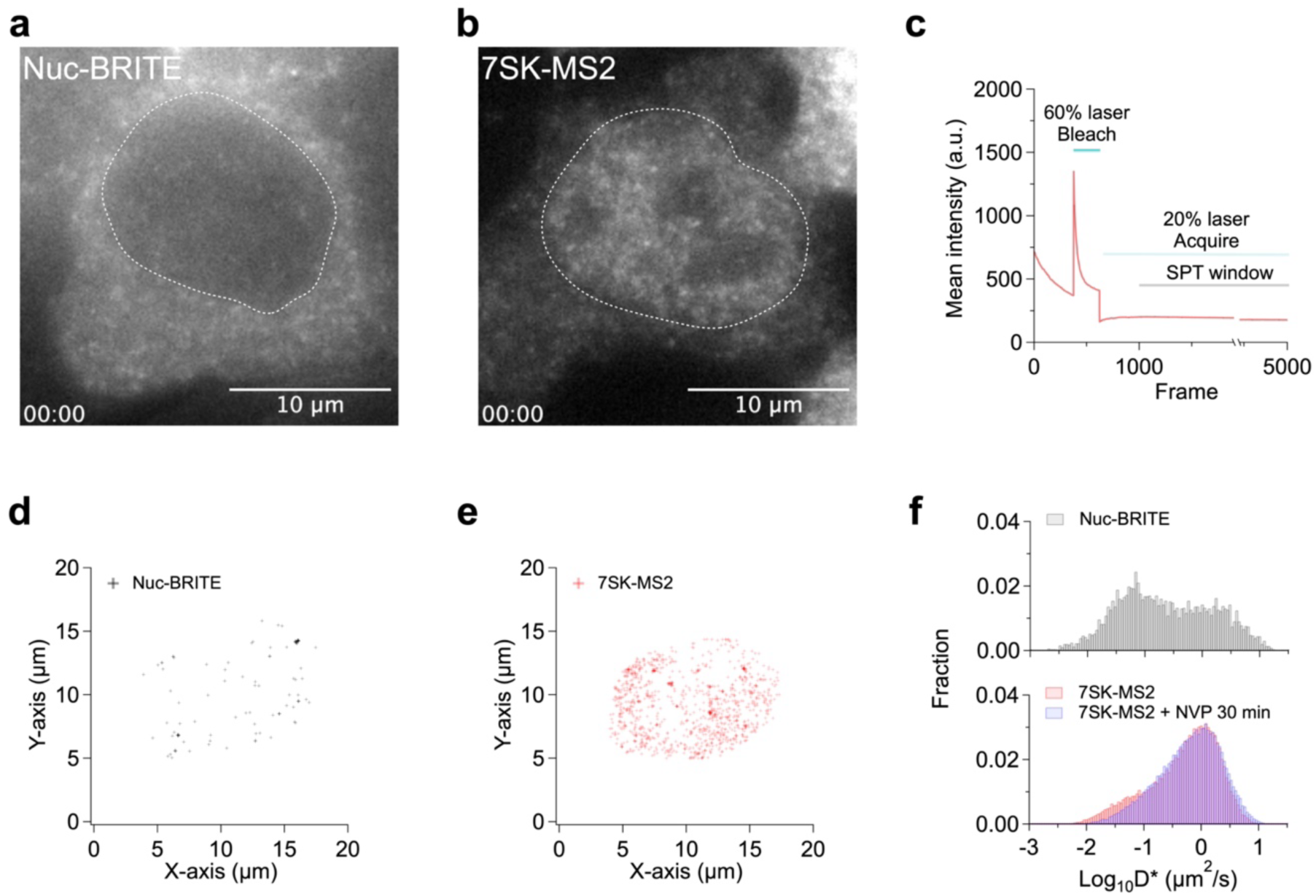
Nuclear-BRITE enables the visualisation of 7SK dynamics using single particle tracking. First frame of SPT acquisition for a) Nuc-BRITE alone and b) 7SK-MS2 showing the observed nuclear signal within the white dashed line. c) Mean intensity profile of SPT acquisition depicting the initial acquisition to mark the nuclear boundary, bleaching with 60% laser power and reduction down to 20% laser power for the subsequent SPT analysis window from frame 1000 – 5000 at a frame rate of 15 ms. Representative mean trajectory localisations for individual d) Nuc-BRITE and e) 7SK-MS2. f) Log_10_D* histograms of Nuc-BRITE, 7SK-MS2 and 7SK-MS2 + NVP-2 samples. Trajectories analysed from 34, (Nuc-BRITE), 130 (7SK-MS2) and 169 (7SK + NVP-2) and 34 (Nuc-BRITE) cells taken from 3 independent replicates.

### Enhanced imaging of nascent transcripts using Nuclear-BRITE

To further quantify the benefits of Nuc-BRITE compared to conventional MS2 nuclear imaging methods we focused on methodologies for imaging nascent transcription^40,53^. To gain sufficient signal-to-noise for long-term imaging, recent studies have inserted MS2 x128 arrays into introns of transcriptionally active genes of interest^52,54,55^. In some scenarios, a targeted gene or intron may be affected by the large size of this tag, as it could influence transcriptional output and splicing efficiency. Furthermore, accurate nascent transcription imaging can be hindered if the transcription site of interest sits in close proximity to the nucleolus, where excess NLS-tdMCP commonly accumulates. Accordingly, we performed a comparison of intron tagging of Hspg2 using either Nuc-BRITE (MS2v7 x24) or an NLS-tdMCP-GFP (MS2 x128) intronic array as described previously^52^ (Fig 5a). The intensity of single Hspg2-MS2 x128 (tdMCP-GFP) molecules was observed to be similar to Hspg2-Nuc-BRITE (tdMCP-tdStayGold-ERT2) molecules (Fig 5b). This is expected due to the increased brightness of tdStayGold compared to GFP^45^ (∼ 3-fold brighter in cells) and in part the differences in affinity and occupancy of the MS2 arrays^20,46,54,56^. Normalising the observed intensities to that of spliced MS2-labelled intron lariats indicates ∼ 1 – 4 nascent RNA molecules at the transcription site using both tagging systems (Fig 5c). Importantly, the signal-to-noise ratios are significantly improved with Nuc-BRITE when compared to MS2 x128 transcripts, despite the MS2v7 x24 array being ∼ 5x smaller (Fig 5d). Short-term imaging using Nuc-BRITE enabled detection of individual splicing events and changes in transcriptional output at single transcript resolution. Additionally, Nuc-BRITE was stable over extended imaging times (8h), allowing for the visualisation of long-term transcriptional bursting behaviour (Supplementary Video 5).

**Fig. 5.**
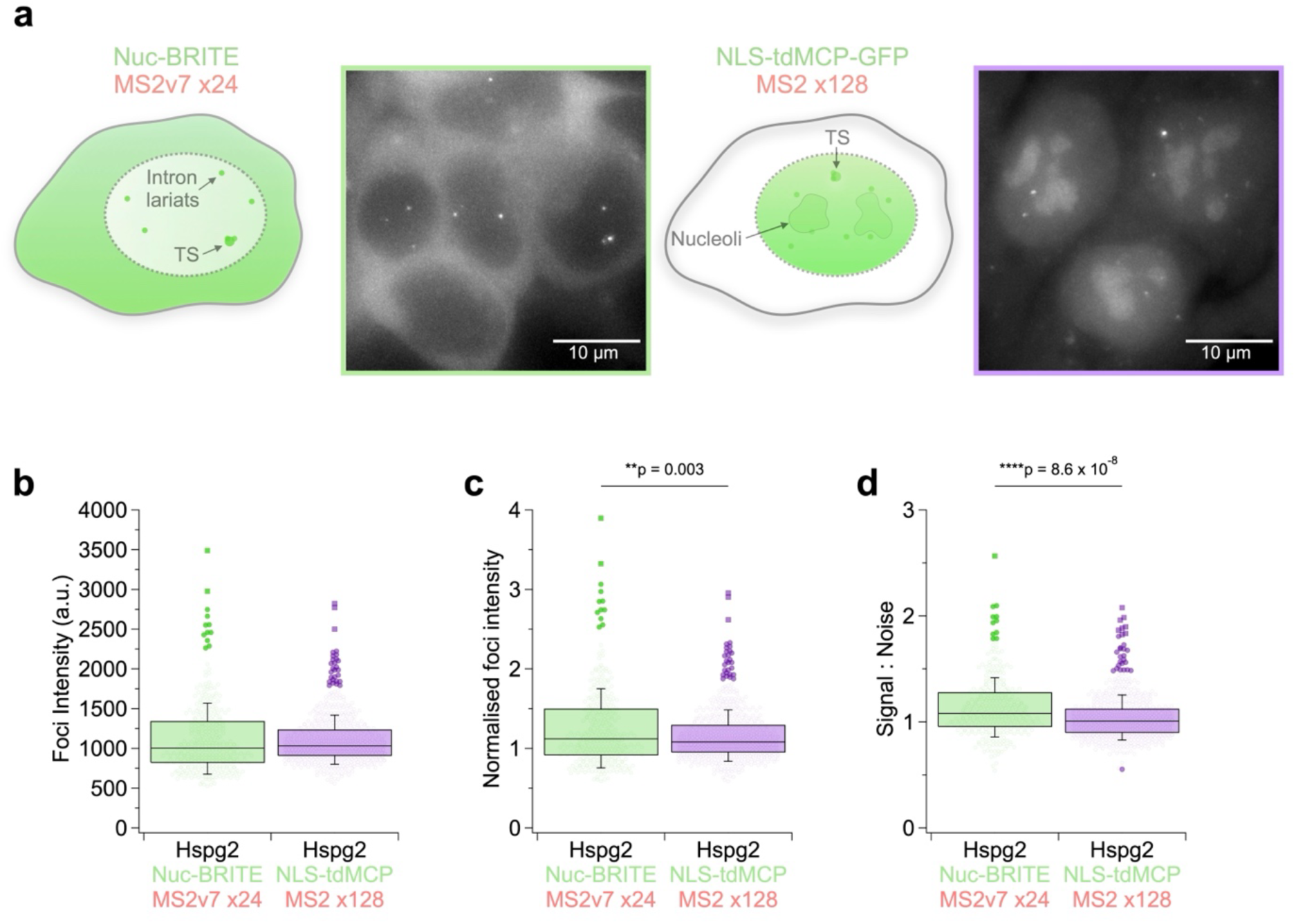
Improving nascent transcript imaging with Nuclear-BRITE. a) Schematics and representative maximum intensity projections of Hspg2 intron 33, labelled with either Nuc-BRITE-MS2v7 x24 or NLS-tdMCP-MS2 x128, showing transcription sites (TS), intron lariats and background nucleolar accumulation in the NLS-tdMCP cells. b) Box plots of Hspg2-Nuc-BRITE and Hspg2-MS2 x128 mean foci intensities, c) normalised foci intensities and d) signal-to-noise ratios. Data acquired from 92 and 106 cells with a total of 329 and 617 foci for Hspg2-Nuc-BRITE and Hspg2-MS2 x128 respectively. All statistical analysis used single tailed students t-tests.

## Discussion

The development of Nuc-BRITE provides a step change in our ability to measure the dynamic behaviour of specific RNAs in the cell nucleus. We demonstrate that Nuc-BRITE is suitable for labelling a broad range of nuclear retained RNAs, both long and short, without visibly affecting their abundance or localisation. The cytoplasmic localisation of Nuc-BRITE fusion proteins does not limit its ability to robustly label highly abundant nuclear RNAs such as Malat-1 and 7SK. A key advance is that we have been able to demonstrate the utility of Nuc-BRITE for imaging endogenously tagged RNAs, contrasting with prior work that validated alternative background-reduced imaging methods using tagged RNAs highly overexpressed from transiently transfected plasmid constructs^23–25,28–30,32–34^. Accordingly, the Nuc-BRITE tagged RNAs in this study are expressed at physiological levels, minimising possible overexpression artifacts. Moreover, homozygous tagging of unique RNAs enables assessment of functionality. A good example of this is Nuc-BRITE tagged 7SK RNA for which loss of function is predicted to have significant detrimental effects on transcription and hence cell viability^57–59^. Further evidence that Nuc-BRITE is minimally disruptive of normal RNA function comes from assessing Polr2a mRNA and 7SK RNA behaviour previously shown to change in response to specific drug treatments^49,50^. Thus, we were able to quantify the dynamic behaviour of both RNAs at single-molecule resolution, either with a MS2v7 x24 array (Polr2a mRNA) or with a single MS2 aptamer within an internal stem-loop (7SK). Upon puromycin treatment, Polr2a mRNA was observed to dissociate from translation dependent accumulations in the cytosol. This effect was rapid (∼ 60 – 120 s) and led to reduced foci intensity and increased mobility as predicted. For 7SK we treated the cells with NVP-2, a CDK-9 inhibitor, which has been shown to reduce chromatin interactions and increase the mobility of 7SK RNP components^50^. We demonstrate equivalent effects on 7SK RNA itself using Nuc-BRITE and SPT.

We show that Nuc-BRITE out competes conventional MS2 imaging in the nucleus when imaging complex nuclear retained transcripts, such as Xist, and even allows for single-molecule detection and tracking at fast frame rates. Building on this capability, we have used Nuc-BRITE labelled Xist RNA to gain important insights into the mechanism of Xist RNA localisation^41^. Of note, Xist and other large nuclear retained lncRNAs can accumulate in high numbers within a limited volume, and consequently fast frame rate live-cell imaging at single-molecule resolution using conventional MS2 approaches is relatively challenging^60^. Nuc-BRITE improves many aspects of single-molecule imaging on RNAs such as Xist, where significant improvements in signal-to-noise ratios, track numbers and track lengths are observed. Additionally, we demonstrate the ability to label MS2 tagged nuclear RNAs in a rapid and tuneable way using Nuc-BRITE-Halo and the addition of Tamoxifen. Alternatively, Nuc-BRITE-tdStayGold allows for robust constitutive labelling of RNAs for wider applicability and long-term imaging experiments. We envisage these two options of Nuc-BRITE constructs will provide valuable new options for controlling the labelling of RNAs of interest.

Finally, we benchmarked Nuc-BRITE as a tool for imaging nascent transcription dynamics using the Hspg2 gene as a model. Quantification of nascent transcription dynamics and transcriptional bursting behaviour is increasingly used as a tool to understand mechanisms of transcriptional regulation^12,40,53,61,62^. Recent studies have been performed successfully using a an MS2×128 array^52,54^, but the relatively large size of this tag increases the possibility that it will interfere with physiological transcription dynamics. We attained equivalent single transcript sensitivity using the smaller MS2v7 x24 array in conjunction with Nuc-BRITE. Moreover, improved signal-to-noise ratios in spite of the MS2v7 x24 array being ∼5 times smaller, affords new opportunities in analysing native gene transcription and splicing dynamics with minimal impact on gene function.

In conclusion, we have developed a novel approach to improve the imaging of endogenous RNAs in the cell nucleus. We have applied this methodology to a broad range of nuclear RNAs of different sizes and expression levels without affecting their function or abundance. We show that imaging previously challenging nuclear RNAs can now be achieved at single-molecule resolution in living cells and at fast frame rates. We believe this methodology will enable the assessment of the dynamic behaviour of previously under-studied nuclear RNAs that play essential roles in gene expression and RNA processing events.

---

**Supplementary Table 1.**
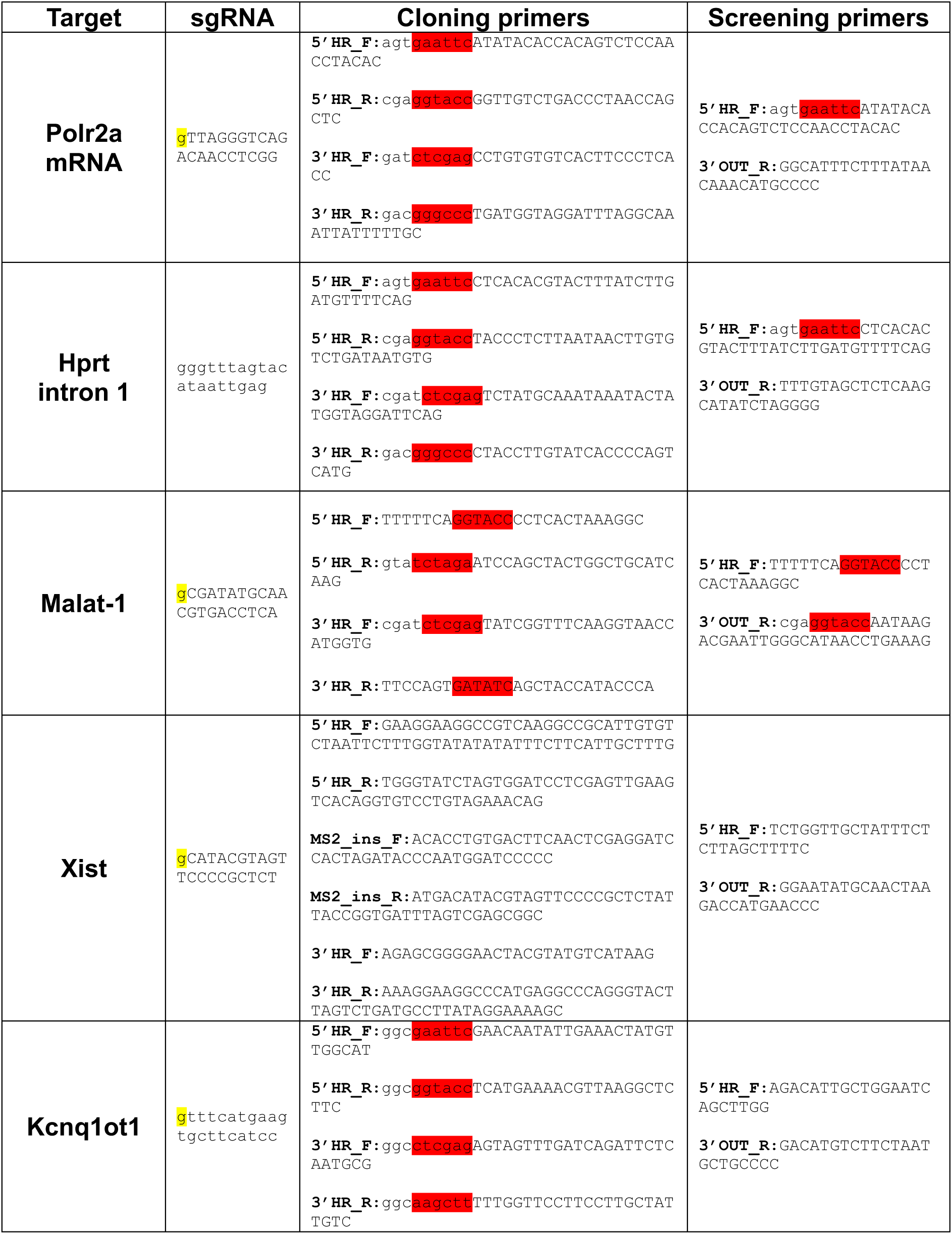
sgRNA sequences used for CRISPR/Cas9 HDR. Yellow highlights additional guanine nucleotide required for efficient PolIII expression. Cloning primers show genic sequences for PCR amplification (capitalised) and restriction enzyme sites (red highlight) used for cloning. Xist cloning primers show primers used for Gibson assembly. Screening primers show 5’ homology region forward primers and 3’ outside homology reverse primers.

**Supplementary Table 2.**
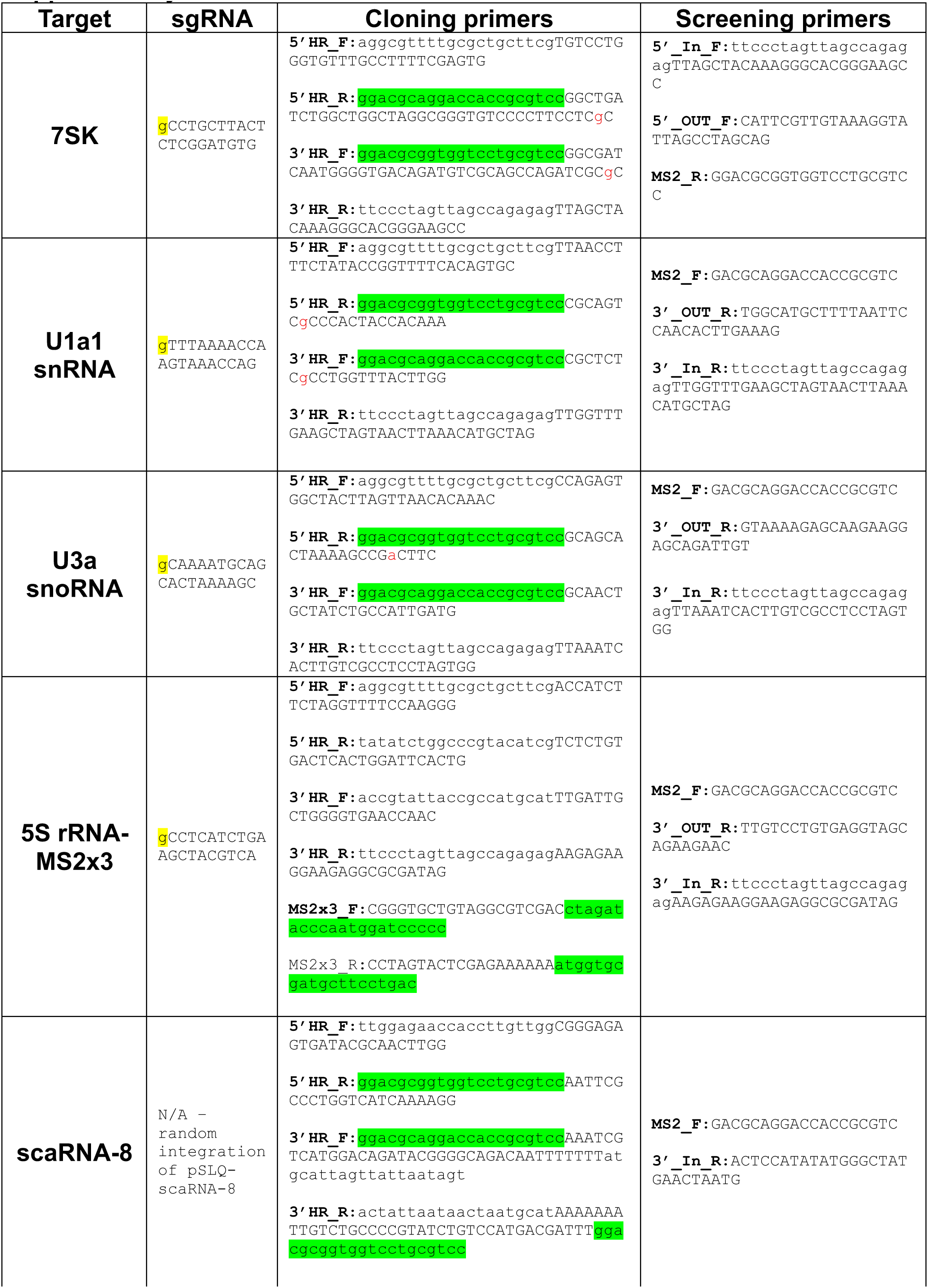
sgRNA sequences used for CRISPR/Cas9 HDR. Yellow highlights additional guanine nucleotide required for efficient PolIII expression. Cloning primers show genic sequences for PCR amplification (capitalised) and Gibson homology arms (lowercase) with MS2 sequences highlighted (green). Screening primers show primers within homology regions (In) and primers outside homology regions (OUT) along with MS2 primers used to detect genomically integrated MS2 sequences.

**Supplementary Table 3.**
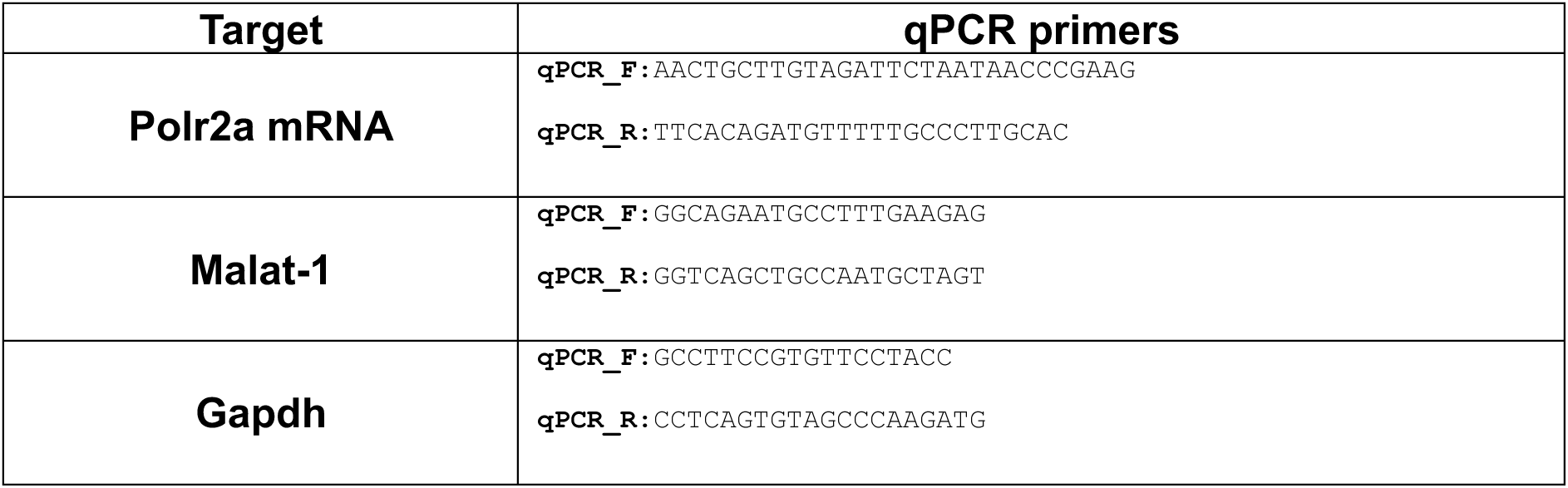
qPCR primers – forward and reverse primers for Polr2a, Malat-1 and Gapdh.

## Acknowledgements

We would like to thank members of the Brockdorff lab, Alexander Borodavka and Stephan Uphoff for critical reading of this manuscript. And funders of grants awarded to N.B. from the Wellcome Trust (215513/Z/19/Z) and UKRI (EP/Y029062/1). Imaging was performed at the Micron Oxford Advanced Bioimaging Unit funded by a Wellcome Trust Strategic Award (091911 and 107457/Z/15/Z).

## Contributions

N.B. and A.D.C conceived experiments. N.B. and A.D.C wrote the manuscript. A.D.C generated and validated cell lines, conducted imaging experiments and image analysis. B.C. generated Kcnq1ot1 cells and conducted live-cell imaging experiments. A.T.S. and R.J.K. provided the Hspg2-MS2×128 cell line, Hspg2 targeting constructs and helped with long-term microscopy and analysis.

## Data, code, and materials availability

All data, code, and materials used in this study are available upon request. MATLAB data analysis pipelines for RNA SPT data are available on github: https://github.com/adamcawte/NucBRITE_MATLAB_Code.git

---

**Supplementary Figure S1.**
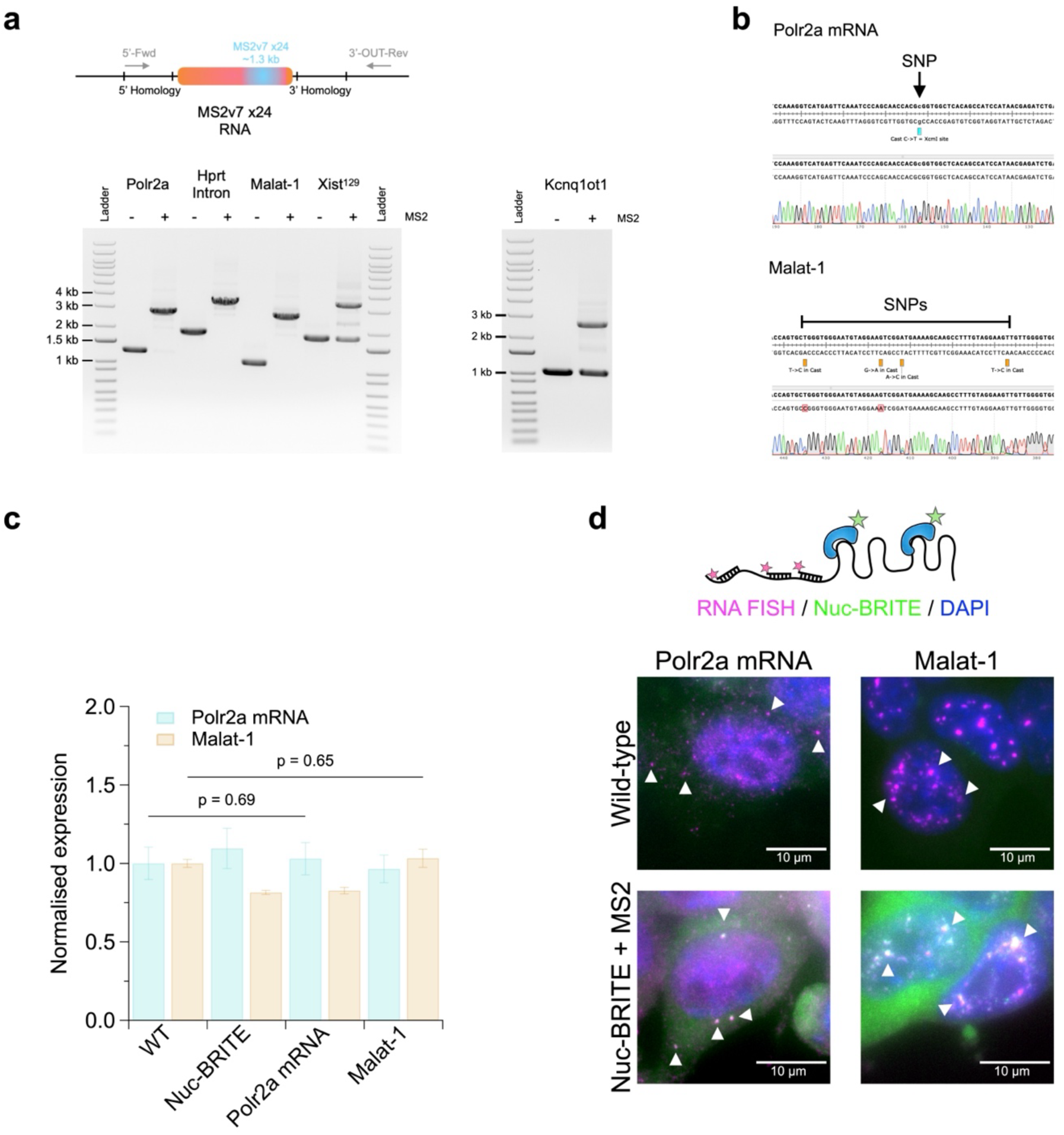
Characterisation of long RNAs tagged with Nuclear-BRITE. a) Schematic and agarose gels for PCR screening of MS2v7 x24 insertion compared to Nuc-BRITE cells lacking MS2 repeats. Forward primers were in the 5’ homology region of the repair template and reverse primers located outside of the homology template to detect MS2v7 x24 insertion (∼1.4 kb) at each endogenous locus. b) Sanger sequencing profiles of MS2v7 x24 tagged Polr2a mRNA and Malat-1 outside of the repair homology showing the presence of 129/Castaneus single nucleotide polymorphisms confirming homozygous insertion of MS2v7 x24. c) Normalised RNA expression of Polr2a mRNA and Malat-1 relative to Gapdh taken from 4 technical replicates with error bars shown as standard deviation around the mean and p-values from a two-tailed students t-test. d) RNA-FISH (magenta) and Nuc-BRITE (green) imaging of Polr2a mRNA and Malat-1 signals in wild-type and MS2v7 x24 tagged Nuc-BRITE cells. Arrows depict colocalization and accumulation of RNA FISH signal with Nuc-BRITE signal.

**Supplementary Figure S2.**
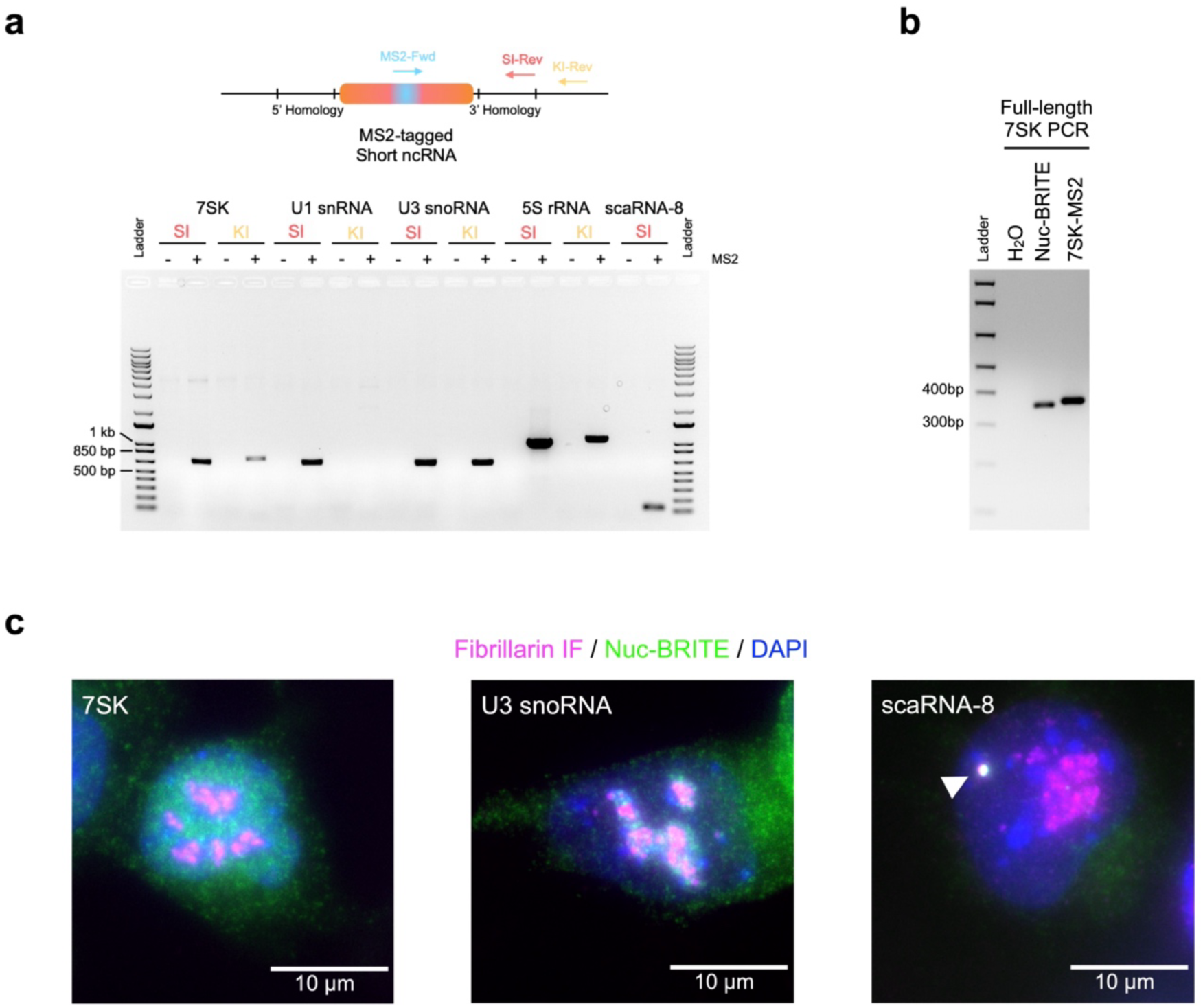
Characterisation of short ncRNAs tagged with Nuclear-BRITE. a) Schematic and agarose gel of PCRs for short ncRNA cell lines to confirm stable integration (SI) or endogenous knock-in (KI). Negative control lanes (-MS2) used Nuc-BRITE genomic DNA lacking MS2 insertion. b) PCR gel of full-length 7SK with or without MS2 incorporation showing homozygous knock-in of internal MS2. c) Maximum intensity projections of Fibrillarin (magenta) and Nuc-BRITE (green) signals for nucleoplasmic 7SK, nucleolar U3 snoRNA and Cajal body specific scaRNA-8, showing each Nuc-BRITE tagged short ncRNA is correctly localising with its target nuclear compartment.

**Supplementary Video. 1**

Fast live-cell super-resolution iSIM imaging of Nuc-BRITE tagged RNAs, Polr2a mRNA, Hprt intron 1, Malat-1, Xist and Kcnq1ot1 lncRNAs. Nuc-BRITE cells without MS2 aptamers present shown for reference. All videos acquired at a 75 ms frame rate and shown in real time except Hprt intron 1 which is a maximum projection of an acquisition taken at a 10 s frame rate to observe RNA splicing events, shown at 5 fps. Scale bars 10 µm.

**Supplementary Video. 2**

Fast live-cell super-resolution iSIM imaging of Nuc-BRITE tagged short ncRNAs, 7SK, U1 snRNA, U3 snoRNA, 5S rRNA and scaRNA-8. Nuc-BRITE cells without MS2 aptamers present shown for reference. All iSIM videos acquired at a 75 ms frame rate and shown in real time. Scale bars 10 µm. Long-term imaging of 5S rRNA nucleolar import taken on an Olympus widefield microscope at a 15 min frame rate shown as a maximum projection of multiple Z-slices at 5 fps.

**Supplementary Video. 3**

Maximum projection of long-term imaging of Polr2a mRNA foci following puromycin treatment. Taken on an iSIM microscope at a 5 sec frame rate over a 10 min time course and multiple Z-sections. Video shown at 5 fps. Fast live-cell super-resolution iSIM imaging of Polr2a mRNA in the absence and presence of puromycin, showing puromycin sensitive cytoplasmic accumulations of multiple Polr2a mRNA molecules (white arrows). Fast imaging acquired at a 75 ms frame rate and shown in real time. Scale bars 10 µm.

**Supplementary Video. 4**

SPT acquisition of Nuc-BRITE and 7SK nuclei. First static frames depict background signal in Nuc-BRITE cells and 7SK specific signal. The following frames show the nuclear boundary (yellow) within which single particles are to be analysed, the subsequent frames show rounds of bleaching prior to single molecule detection. All foci detected in TrackMate are shown, but only later frames (1000 – 5000) are analysed depicted as coloured circles and traces, excluded foci shown in dark grey. SPT acquired at a 15 ms frame rate and shown in real time. Timestamp depicts elapsed video time in a minutes : seconds format. Scale bars 10 µm.

**Supplementary Video. 5**

Maximum projections time series of long-term imaging of nascent Hspg2 foci for either 1 h at 1 min frame rate, showing single transcript sensitivity and intron splicing – as quantified in the adjacent graph, or for 8h at a 4 min frame rate showing long-term transcriptional bursting behaviour. Scale bars 10 µm and time shown as hour : min.

